# The role of stochastic sequestration dynamics for intrinsic noise filtering in signaling network motifs

**DOI:** 10.1101/278929

**Authors:** Debdas Paul, Nicole Radde

## Abstract

The relation between design principles of signaling network motifs and their robustness against intrinsic noise still remains illusive. In this work we investigate the role of cascading for coping with intrinsic noise due to stochasticity in molecular reactions. We use stochastic approaches to quantify fluctuations in the terminal kinase of phosphorylation-dephosphorylation cascade motifs and demonstrate that cascading highly affects these fluctuations. We show that this purely stochastic effect can be explained by time-varying sequestration of upstream kinase molecules. In particular, we discuss conditions on time scales and parameter regimes which lead to a reduction of output fluctuations. Our results are put into biological context by adapting rate parameters of our modeling approach to biologically feasible ranges for general binding-unbinding and phosphorylation-dephosphorylation mechanisms. Overall, this study reveals a novel role of stochastic sequestration for dynamic noise filtering in signaling cascade motifs.

## 1 Introduction

Intracellular signaling pathways are known to function reliably and are robust to all kinds of perturbations and noise. They are often controlled by a complex regulation structure involving network motifs such as positive and negative feedback and feed-forward loops, and a multitude of examples demonstrate that network architecture is tightly related to a robust functioning (Batchelor and Goulian, 2003; Blüthgen and Legewie, 2013; Caicedo-Casso et al., 2015). In particular, the interplay between interlinked positive and negative feedback is often crucial for a precise tuning of these pathways and thus to enable a reliable functioning. Examples include robustness due to negative feedback such as in MAPK signaling (Fritsche-Guenther et al., 2011; Clodong et al., 2007), robustness of oscillations via coupling of feedback loops (Cheng et al., 2001; Wagner, 2005), or fine tuning of thresholds in bistable systems via nested negative feedback mechanisms (Justman et al., 2009). Another well-studied kind of motif that is related to thresholding and switching behavior is multisite phosphorylation (Gunawardena, 2005; Whitmarsh and Davis, 2016) and, as a special case, cascades of phosphorylation-dephosphorylation (PD) cycles as they can be found in several pathways such as the MAPK or AKT signaling pathway (Angeli et al., 2004). Cascading prolonges the duration of signal propagation and hence acts as a low-pass filter which filters out high frequencies (Paul and Radde, 2016). Moreover, even in signaling cascades without explicit feedback, where one PD cycle activates the next PD cycle downstream, information does not only propagate from upstream to downstream molecules, but also in the opposite direction, an effect that is called retroactivity (Del Vecchio et al., 2008; Del Vecchio and Sontag, 2009; Ventura et al., 2008, 2009, 2010). The presence of retroactivity has been verified theoretically and experimentally for example in the MAPK signaling pathway (Kim et al., 2011b, 2010). From a clinical perspective, retroactivity is of particular interest because it facilitates off-target effects of kinase inhibitors, a class of extremely effective anti-cancer agents (Wynn et al., 2011). Moreover, retroactivity acts as an implicit feedback and therefore plays an important role in the dysregulation of signaling networks (Wynn et al., 2011). Therefore retroactivity, in general, has been considered as a design trade-off which must be minimized or attenuated as far as possible to facilitate unidirectional signal propagation in signaling pathways (Shah and Vecchio, 2017). However, in this work we reveal a new effect of retroactivity, which can contribute to reducing intrinsic noise and fluctuations in the activity of the terminal kinase of a double PD cycle cascade. We compare models of a simple double PD cycle motif (Fig. 1, model A) and a cascade of two of such motifs (Fig. 1, model B) in terms of stochastic simulations and variations in the activity of the terminal kinase. We observe that in model B time-varying sequestration of ppX plays a significant role in regulating the extend of intrinsic noise in ppY. This is a purely stochastic phenomenon and has no counterpart in deterministic regimes. Therefore, we named it *dynamic sequestration*. We put our results into a realistic biological context by using biologically feasible ranges of rate parameters. Overall, our results elucidate the role of stochastic sequestration dynamics for the regulation of output variability in double PD cycle cascade motifs without explicit feedback control.

**Fig. 1.**
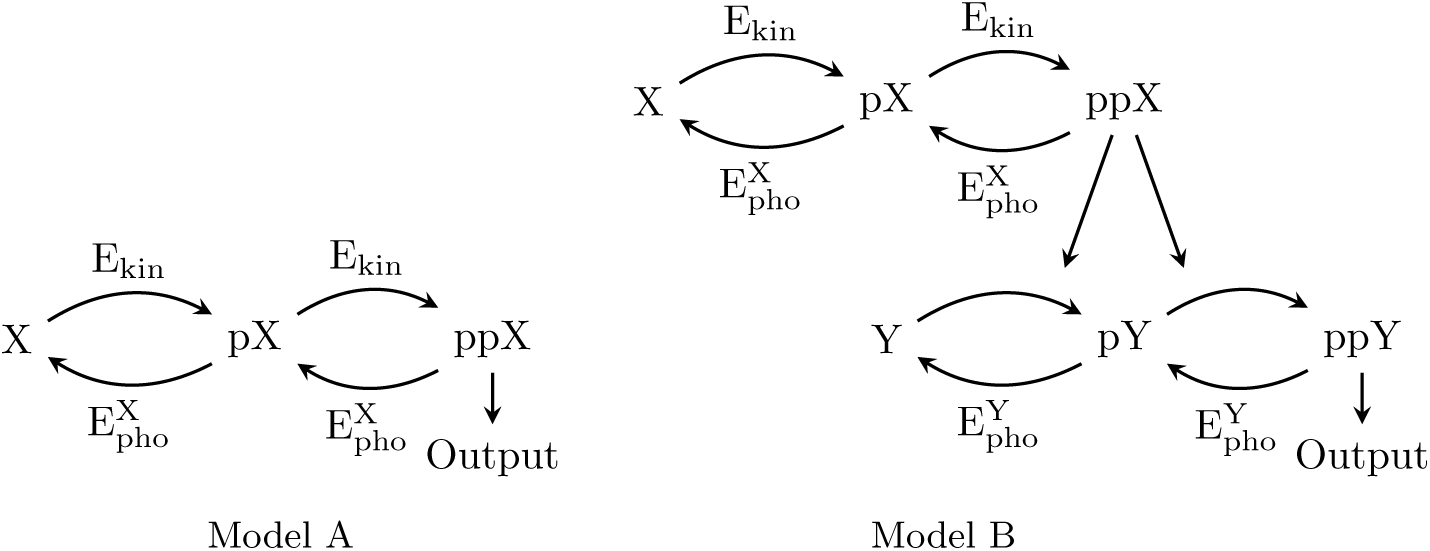
Different cascade motifs of phosphorylation-dephosphoryation cycles. (*Left*) A double PD cycle motif (model A) and (*Right*) a cascade of two of such motifs (model B) are compared in this study. X and Y are different proteins, pX and ppX denote single and double phosphorylated forms of protein X, and the same notation holds for the protein Y. Phosphorylation and dephosphorylation are triggered by kinase and phosphatase molecules, E_kin_ and 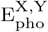, respectively.

## 2. Results

We use Gillespie’s direct stochastic simulation algorithm to generate stochastic sample paths for models A and B. Therefore, we model phosphorylation and dephosphorylation by kinase and phosphatase molecules by adopting the two step enzyme-substrate kinetics to describe the corresponding biochemical reactions (see Section 4.1 for more details). The notations that we use throughout the paper are listed in Table 1. Expectation values are estimated via the generation of many sample paths and Monte Carlo integration and refer to an equilibrium state in which the expectation is time-invariant. Variability in the outputs is mainly compared via coefficients of variation (CV), which is, unlike the standard deviation or the variance, a dimensionless quantity and is appropriate for a comparison of variables with different units or with large differences in their mean values.

**Table 1.** Notation.

| Notation | Meaning |
| --- | --- |
| $\text{ppX}^t$ | Total number (including complexes) of doubly phosphorylated X protein molecules |
| $Y^t + \text{pY}^t$ | Total number of not doubly phosphorylated Y protein molecules |
| $Y^T$ | Total number Y protein molecules, considered as a conserved quantity |
| $\mathbf{E}[\overline{\text{ppX}}], \mathbf{E}[\overline{\text{ppY}}]$ | Expected number of unbound ppX and ppY molecules in steady state |
| $\mathbf{E}[\overline{\text{ppX}}^t]$ | Expected number of total ppX molecules in steady state |
| $k_{\text{on}}, k_{\text{off}}, \text{ and } k_{\text{cat}}$ | Binding, unbinding and catalytic rate constants for a two step enzyme-substrate kinetics (see Section 4.1 for more details) |
| $B_{k^X=k^Y}$ | $k_{\text{on}}, k_{\text{off}}$ and $k_{\text{cat}}$ values of the X protein and the Y protein modules are pairwise equal |
| $B_{k_{\{\text{off}, \text{cat}\}}^Y = r * k_{\{\text{off}, \text{cat}\}}^X}$ | $k_{\text{off}}$ and $k_{\text{cat}}$ of the Y protein module are $r \in \mathbb{R}_{>0}$ times that of the X protein module |
| $c_v^{ss}$ | Coefficient of variation of model output (ppX or ppY, respectively) in steady state |
| $c_v^M$ | Coefficient of variation of steady state output of model M |
| $\rho$ | Pearson's correlation coefficient |
| $r_s$ | Spearman's correlation coefficient |

### 2.1. Retroactivity via dynamic sequestration reduces output variability in cascades of PD cycles

We compared CVs of both model outputs by first using the same set of parameters for the X protein and the Y protein module exemplarily for three different numbers of kinase molecules (Fig. 2). An interesting observation is that for all values of E_kin_, the value of 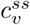 for **E**[ppY] of model 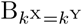 are found to be lower than that of model A. This observation appears counterintuitive at first glance since adding a further stochastic module might increase rather than decrease intrinsic noise, as stated e.g. in Klipp and Liebermeister (2006). Since both modules are identical, the reduction must be caused by the different input signals those modules face. While the X protein module faces E_kin_ as a constant input, since E_kin_ is assumed a conserved quantity, the Y protein module is subject to ppX as input, which is a random variable that changes over time. The reduction in the CV could either be caused by differences in E_kin_ and 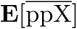, or by the fact that ppX is a time-dependent variable.

**Fig. 2.**
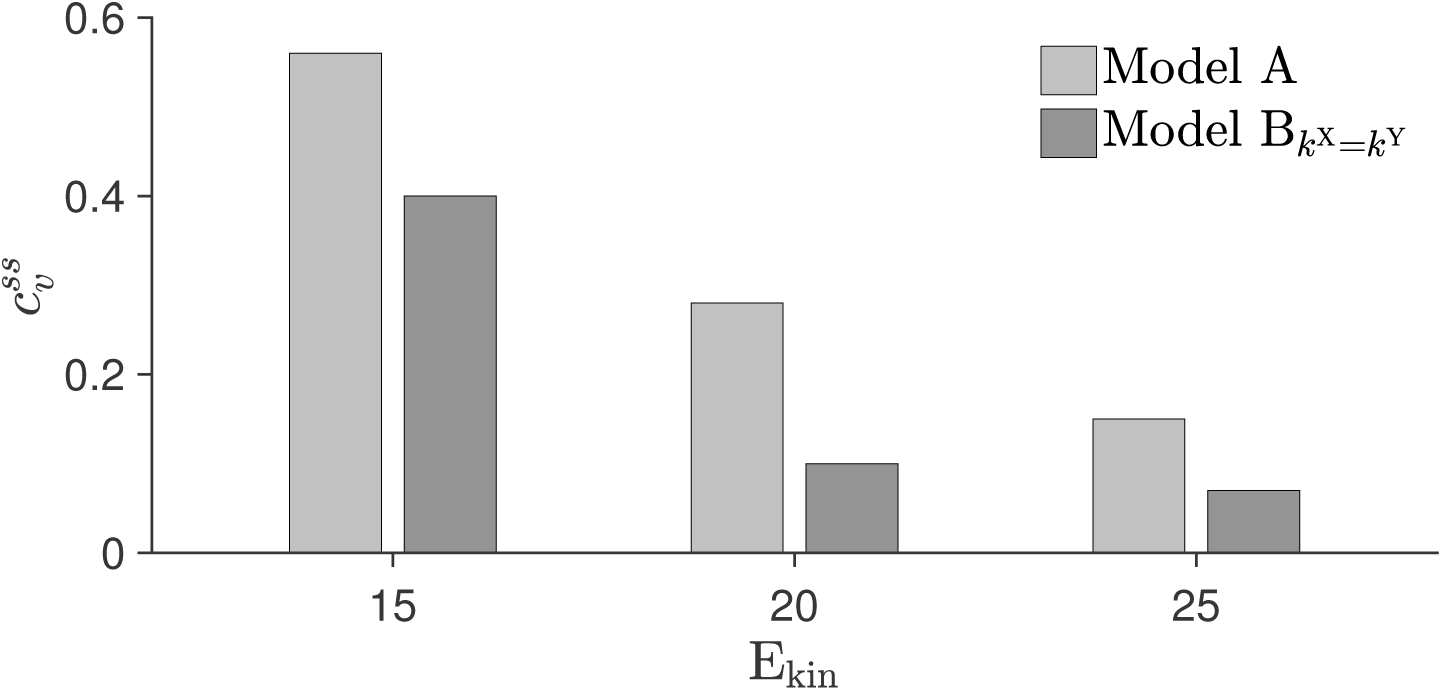
The coefficient of variation 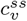 decreases for cascaded architectures. For models A and B, 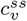are computed for 15, 20 and 25 E_kin_ molecules from an ensemble average of 1000 SSA realizations. Parameter values are summarized in Table 2.

To differentiate between both causes, we recorded the values of 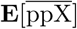 for a range of kinase molecules, as illustrated in Fig. 3 (*left*). While 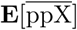 is lower than E_kin_ if E_kin_ is set to 20 molecules or below, it exceeds E_kin_ when E_kin_ is higher than 21 molecules. For E_kin_ = 21 molecules both values are almost identical, suggesting that the reduction in the CV we observed in Fig. 2 is caused by stochastic fluctuations in ppX. Indeed, a comparison of the CVs of model B and of the Y protein module when facing the constant input 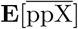 for E_kin_ = 21 molecules (Fig. 3 (*right*)), shows that the coefficient of variation is much larger in the latter case, which supports our suggestion that the stochastic dynamics in ppX is responsible for the reduction in the CV of model B.

**Fig. 3.**
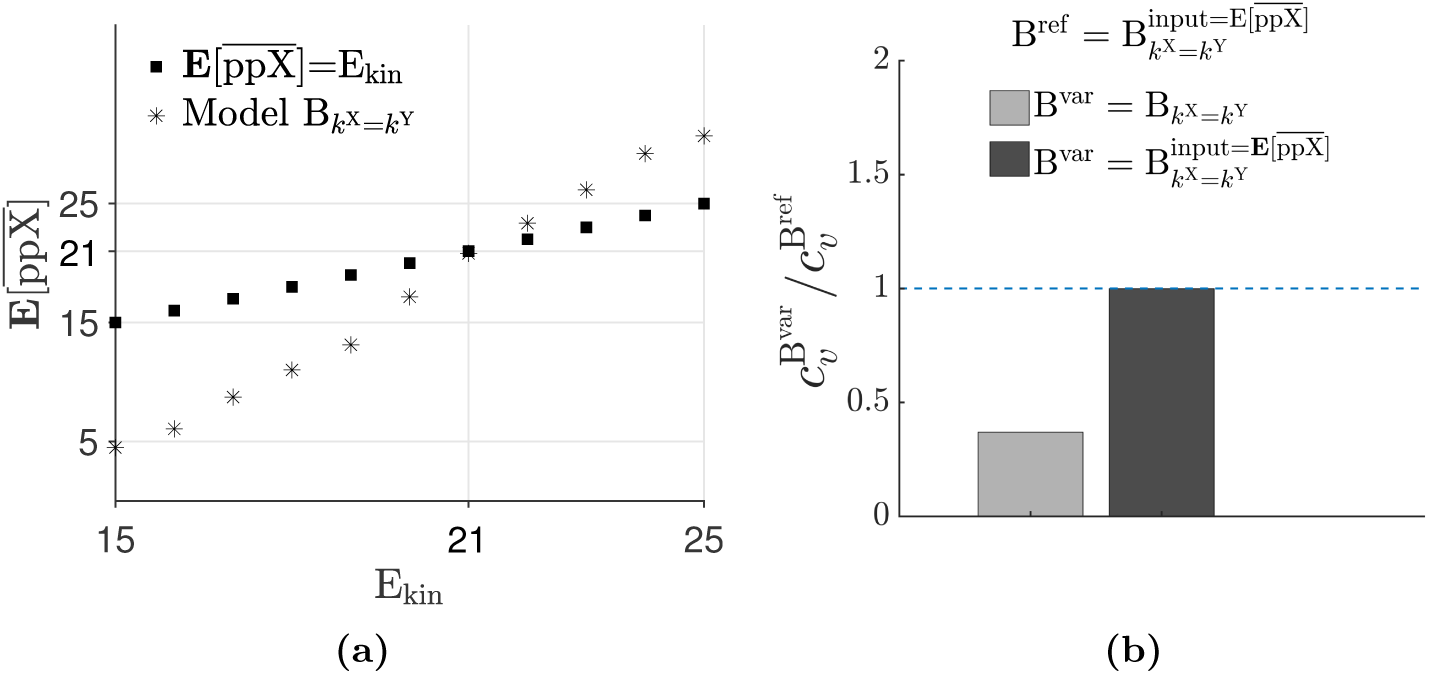
Constant input to the Y protein module does not contribute to the reduction in the value of 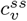 of 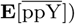. (a) 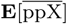 is quantified across a range of E_kin_ molecules. At E_kin_ = 21 molecules, the amount of free ppX takes almost the same value, providing a good reference for further comparisons. (b) Taking E_kin_ = 21 molecules, *c*^*ss*^ values are compared between model B and the case where the Y protein module of model B is fed with a constant input which is set to the expected number of free ppX molecules 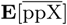. Table 2 summarizes the rest of the parameter values.

Since the underlying stochastic process is characterized by sequestration of ppX by the Y protein module, in a next step we increased the sequestration rate by setting 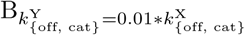, which causes ppX-Y and ppX-pY complexes to accumulate due to slow unbinding and catalytic rates. This further decreased the values of 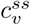, as shown in Fig. 4a, confirming that the output variability is indeed controlled by stochastic dynamic sequestration of ppX. With the same analysis as used before, we explicitly excluded differences in mean values of the input of the Y protein module to be responsible for this further reduction of the 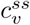 (Figs 4b and 4c).

**Fig. 4.**
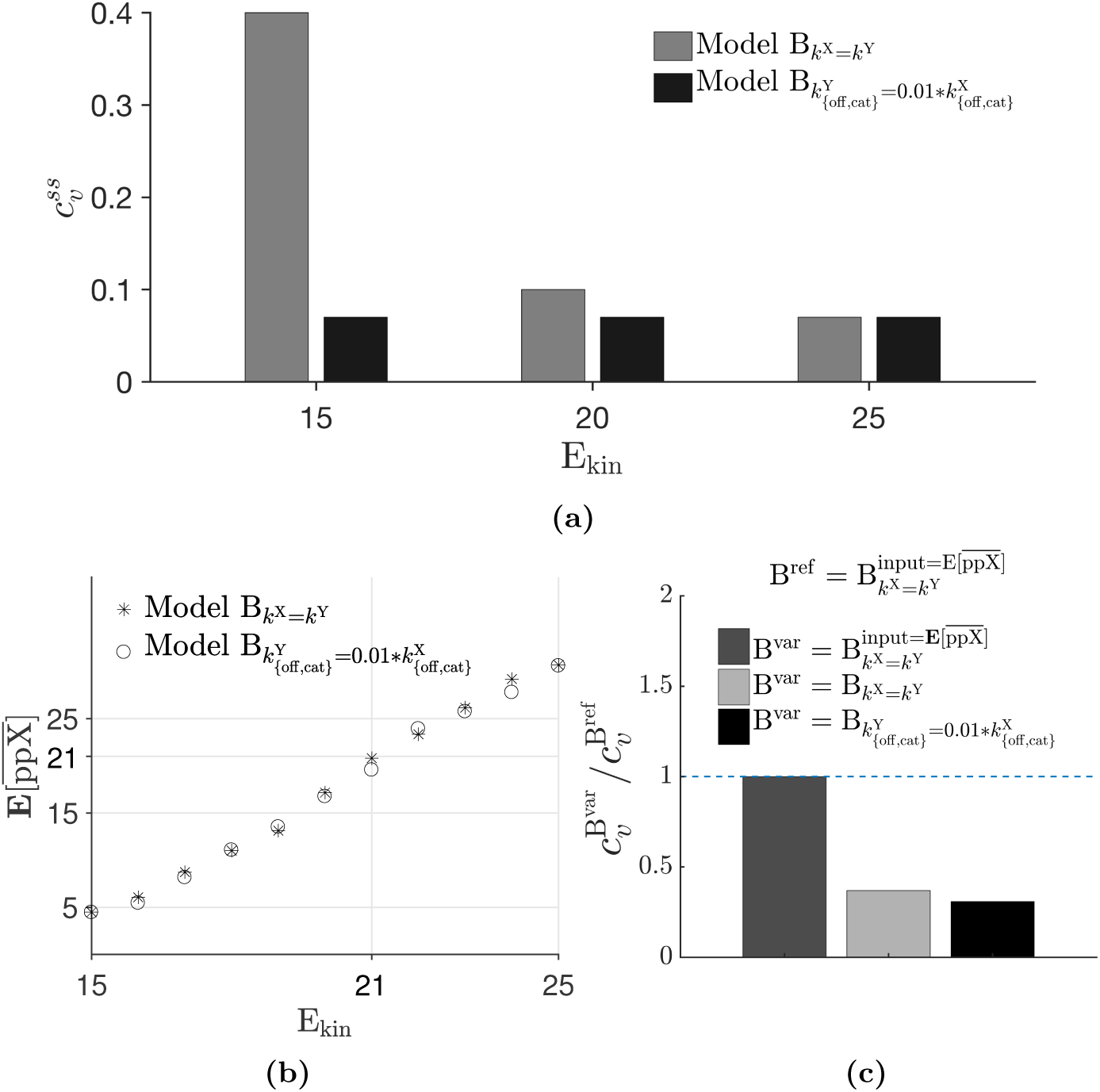
The coefficient of variation 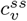 of the cascaded architecture further decreases for slow timescales of the unbinding and catalytic reactions. (a) A comparison of 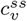 for model B and model B with 100 times smaller *k*_off_ and *k*_cat_ values for the Y protein module, (b)-(c) the same analysis as in Fig. 3 was performed to ensure that this further reduction is indeed primarily caused by the increased sequestration rate of ppX. Table 2 summarizes the rest of the parameter values.

### 2.2. Sensing the downstream module via stochastic dynamic retroactivity

As discussed in the introduction, retroactivity is a well known effect in modular signaling motifs. Here, the phosphorylation state of the X protein molecule is regulated by the downstream molecule Y. Such an effect has experimentally been observed for example in Kim et al. (2011a), where the amount of doubly phosphorylated ERK in the MAPK signaling pathway has been shown to correlate with the number of ERK substrate molecules. The more substrate molecules are available, the more ppERK is sequestered by binding to these substrates. Since those ppERK molecules are temporarily not available for the phosphatase, this sequestration affects the ratio of phosphorylated and unphosphorylated ERK towards higher phosphorylation levels. In this way, ERK can adapt its activity to the number of available substrates.

We investigated if a similar effect is also visible in our model setup and the X system is able to adapt to the state of the Y system. Therefore, we mimicked experiments in Kim et al. (2011a) by calculating the correlation coefficient *r*_*s*_ between 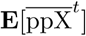 and Y^*T*^. Results are shown in Fig. 5.

**Fig. 5.**
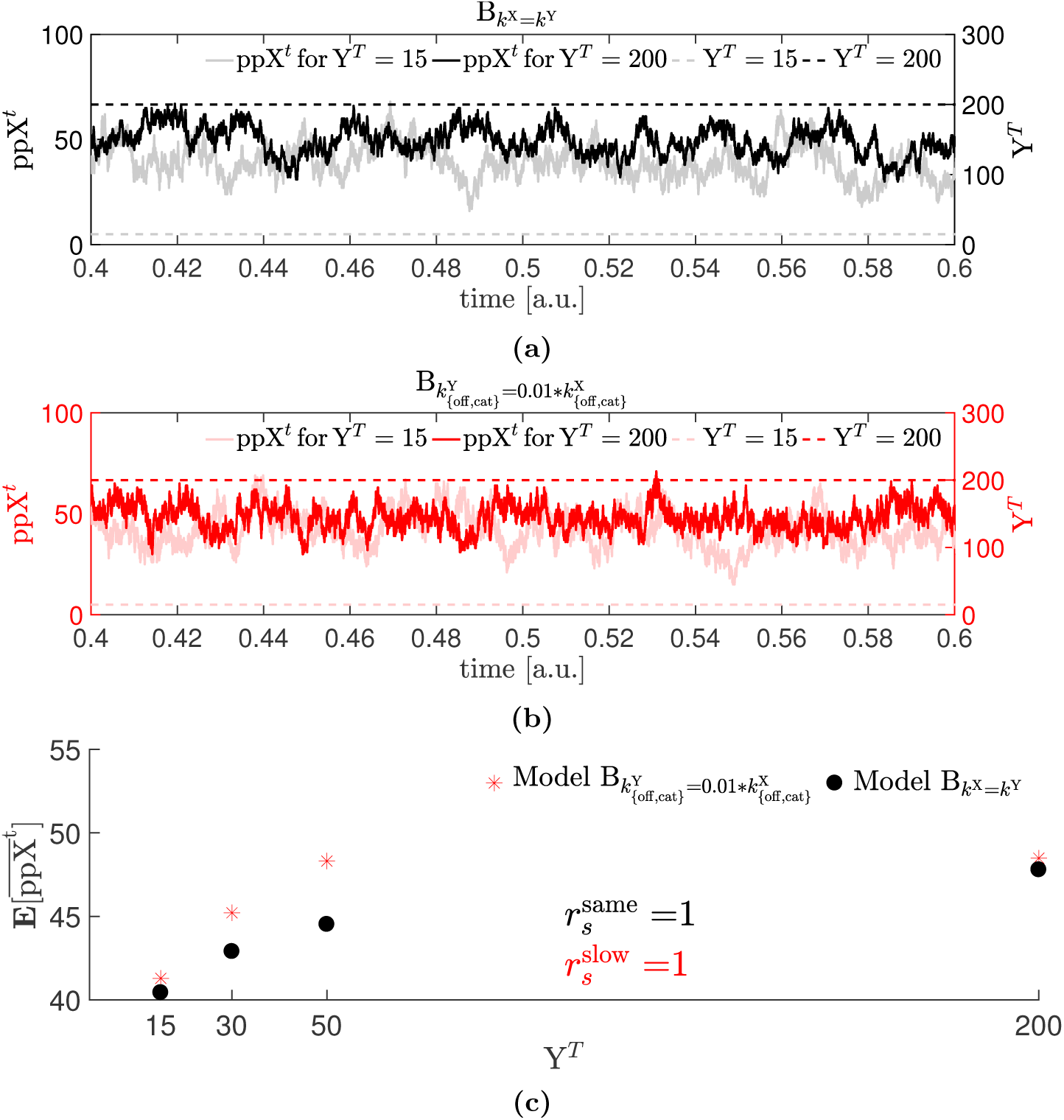
The X protein module senses Y^*T*^ via adaptation of the sequestration rate. Sample paths for ppX^*t*^ in simulations with *Y* ^*T*^ = 15 and *Y* ^*T*^ = 200 molecules for models 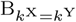 and 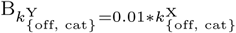 (b). Both settings result in a perfect rank correlation between Y^*T*^and 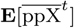.

Shown are representative sample paths of ppX^*t*^ for both 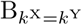 (Fig. 5a) and 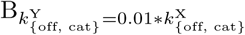 (Fig. 5b) for *Y*^*T*^ = 15 (light gray and red lines, respectively) and Y^*T*^ = 200 molecules (dark gray and red lines, respectively). Thus, model B is able to capture this experimentally observed behavior, confirming that the X molecule senses the needs of the Y molecule via sequestration and adapts to the state of the Y system.

However, in our simulation scenarios in Fig. 4, Y^*T*^ is a conserved quantity. Thus, retroactivity cannot be directly explained by variations in Y^*T*^. The quantity that fluctuates stochastically in our setting is the number of Y molecules that take part in the sequestration, i.e. those Y molecules Y^*t*^+ pY^*t*^ that are not fully phosphorylated. We anticipate that the X protein module is able to sense these fluctuations and to adapt accordingly such that less fully phosphorylated Y molecules trigger a shift in the X protein module towards ppX, resulting in a dynamic correlation between ppX^*t*^ and Y^*t*^+ pY^*t*^. We furthermore anticipate that the strength of these dynamic correlations is highly dependent on the dynamic range in which the whole system operates. For example, the X protein module must be fast enough compared to fluctuations in the Y protein module in order to be able to react to those changes. If this is not the case, the X protein module is too slow to adapt and fluctuations are averaged out.

Therefore, we analyzed the dynamic correlations between ppX^*t*^ and Y^*t*^+ pY^*t*^ directly of the sample paths in different settings. Results are illustrated in Figure 6, which are arranged in a similar manner as in Fig. 5. Figs 6a and 6b show representative sample paths for 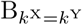 and 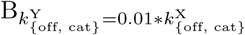 respectively. Simulations were performed with a total number of Y^*T*^ = 100 molecules. Respective distributions of correlation coefficients *ρ* obtained via 1000 simulation runs are shown in Fig. 6c. Both settings show correlations that are significantly different from zero. In the first scenario, time scales for the dynamics in the X and the Y protein modules are comparable, leading to a time delay in the reaction of the X protein module to changes in the Y protein module, which results in an overall negative correlation between ppX^*t*^ and Y^*t*^+ pY^*t*^. In the second scenario, where the Y protein module has a much slower dynamics, the X protein module is able to follow changes in the phosphorylation state of the Y protein module instantaneously, and the correlation is positive instead.

**Fig. 6.**
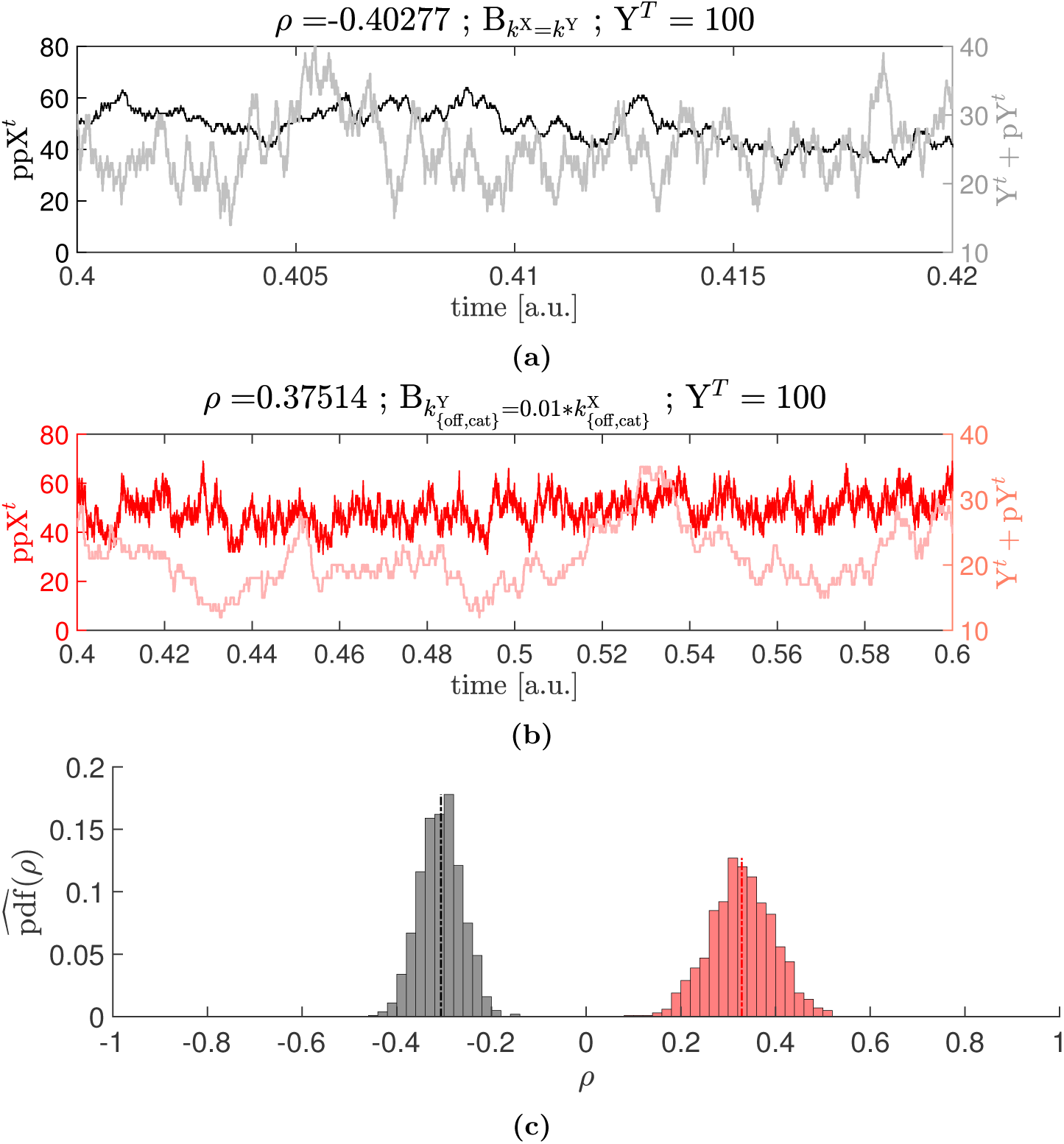
Stochastic sequestration dynamics causes a reduction in output variability of model B. Sample paths of ppX^*t*^ and Y^*t*^+ pY^*t*^ for 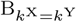 (a) and 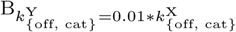 for Y^*T*^ = 100 molecules (b). (c) Distributions of correlation coefficients *ρ* for both settings that have been inferred via 1000 SSA simulations. Parameter settings are listed in Table 2. Param values: Y^*T*^ = 100, E_kin_ = 21 molecules, *μ*_1_ = *-*0.3068, *σ*_1_ = 0.0447 (black) *μ*_2_ = 0.3287, *σ*_2_ = 0.0672 (red)

As expected, correlations become smaller with decreasing number of Y^*T*^ molecules, as exemplarily shown in Fig. A.9 in the appendix, where we have used Y^*T*^ = 15 molecules.

Overall, our analysis shows that stochastic sequestration dynamics affects variability in the activity state of the downstream protein of cascades of double PD cycle motifs. Of note is that this form of retroactivity can only be observed in a stochastic environment and has no counterpart in a deterministic regime.

### 2.3. Dynamic sequestration in biological systems

In order to put our results into biological context, we decided to set model parameters within biologically feasible ranges where applicable. For this purpose, we adopted our parameters by using values recorded in Table 1 of Dhanan-janeyulu et al. (2012). These values have been inferred from experiments on the Ras/MEK/ERK signaling cascade in mammalian cells, as described in Fujioka et al. (2006). Resulting parameter values are listed in Table 3. Using these values, we performed the same analysis as in Fig. 2. Results are recorded in Fig. 7.

**Fig. 7.**
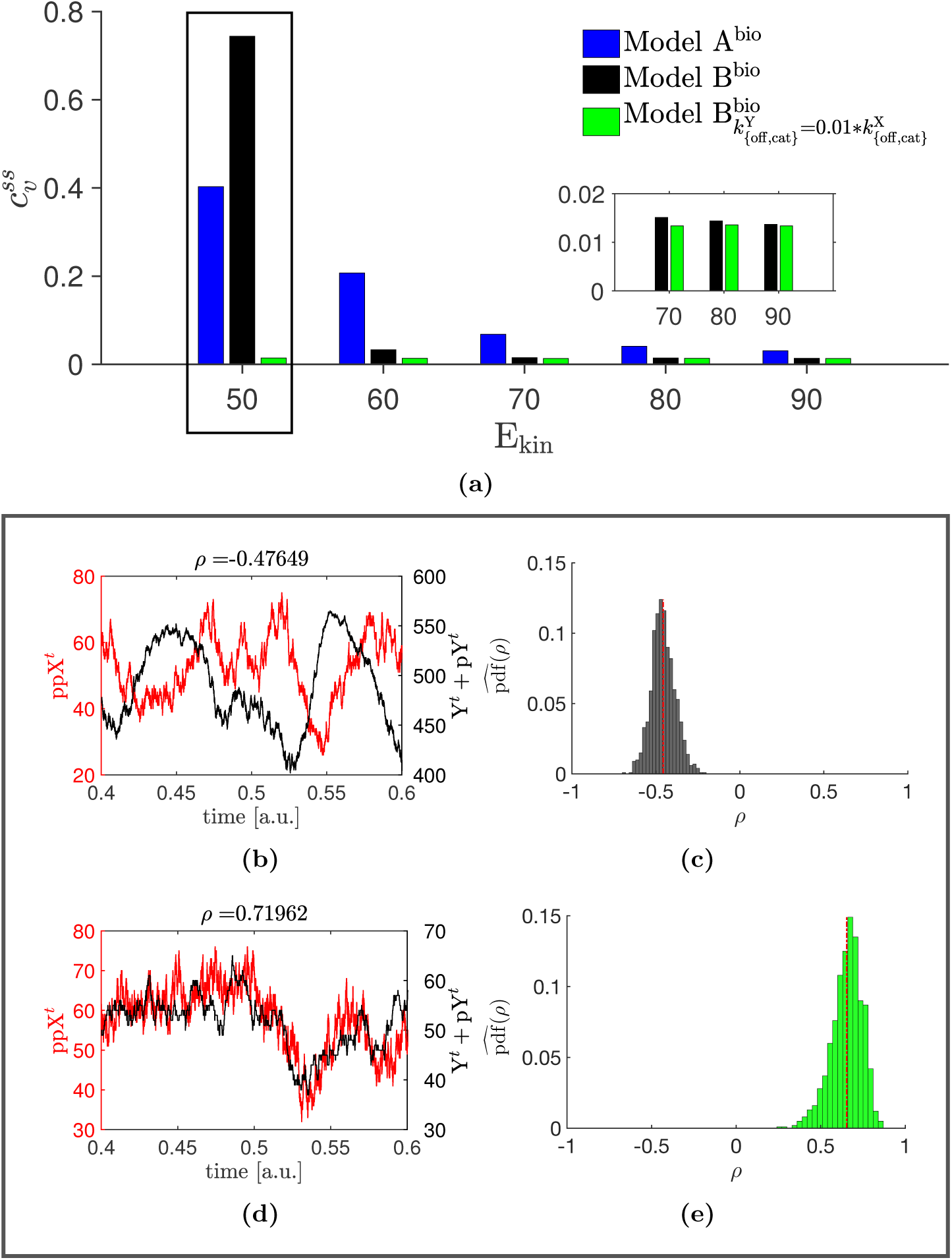
Stochastic sequestration dynamics in a biological context. (a) Same analysis as in Fig. 2. Different model variants are compared in terms of their coefficients of variations 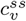. (b)-(e) Sample path and correlation analysis for E_kin_ = 50 molecules for model B^bio^ (b-c) and model 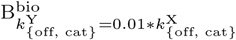 (d-e). Parameters are recorded in Table 3.

While Fig. 7a clearly shows the reduction in the coefficient of variation 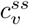 from model A (denoted A^bio^) to model B (B^bio^) and model B with reduced rate constants for the Y protein module (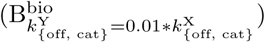) for the range E_kin_ = 60 *-* 90 kinase molecules, model B has a considerably higher output variability for the case E_kin_ = 50 molecules as compared to model A.

To explain this behavior, we did a sensitivity analysis via dose response curves, in which we analyzed model outputs with respect to different input values. Results are shown in Fig. 8 for the ‘non-biological’ context (left column) and the biological context (right column). The left column shows that 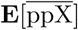 and 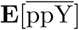 are both in a highly dynami<u>c ran</u>ge for E_kin_ = [15 *-* 25] (indicated by shaded regions). For this regime, 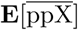 approximately spans a range between 5 and 50 molecules, the respective range for 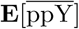 is between 30 and 60 molecules. Thus, both variables are highly sensitive to variations in the input, although 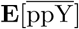 to a lesser extent. For 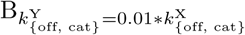, the 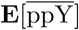 curve increases much faster and is in saturation at about 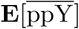 molecules already at E_kin_ = 10.

**Fig. 8.**
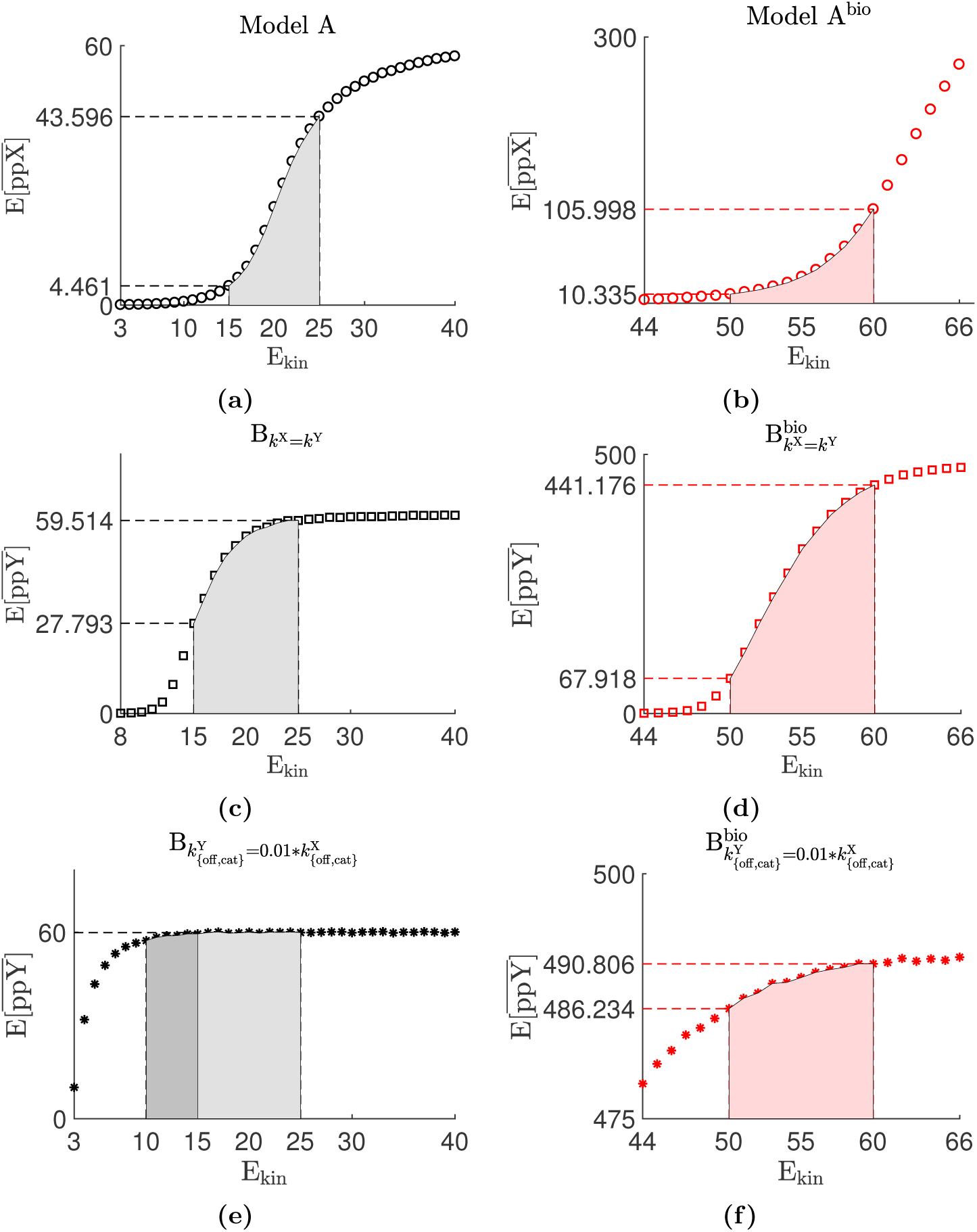
Dose-repsonse curves for model outputs. Expectation values for model outputs as functions of E_kin_ for the ‘non-biological’ context (left) and the biological context (right).

Thus, 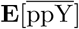 is extremely insensitive to variations in E_kin_ and to stochastic fluctuations in ppX. The results for simulations in the biological context are illustrated on the right. Especially for E_kin_ = 50 molecules, 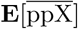 is still at an extremely low value with a small sensitivity, while 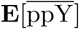 has just reached the start of its dynamic range and thus shows a high sensitivity with respect to variations in E_kin_, which explains the increase of the coefficient of variation in Figure 7a. As before, for 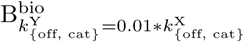, 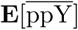 has reached saturation for the range of E_kin_ values that are considered here and hence shows extremely low sensitivities and low coefficients of variation. For higher E_kin_ values, 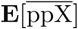 rapidly comes into its dynamic range, while 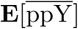 is already almost saturated for E_kin_ = 60 molecules, explaining the immense decrease in the coefficient of variation in model B^bio^ from E_kin_ = 50 to E_kin_ = 60 molecules.

Taken together, this analysis shows that the dynamic range in which the system operates is crucial for the effect of cascading on the output variation and also highly influences stochastic sequestration dynamics.

## 3. Discussion and conclusions

In this study we compared a double PD cycle model (model A) and a cascade of two of such models (model B) with respect to stochastic variations in the activity of the downstream protein. Our analysis revealed an ambivalent role of stochastic sequestration dynamics for the regulation of the Y protein module variability, here measured in terms of coefficients of variation in doubly phosphorylated Y, ppY. Sequestration of doubly phosphorylated X, ppX, by the Y protein module constitutes a kind of retroactivity. Via sequestration, the X protein module senses and reacts to the state of the Y protein module, and hence information is propagated from the downstream to the upstream molecules in these protein cascades. This effect causes a correlation in the sample paths of ppX^*t*^ and those molecules of the Y system that are not fully phosphorylated, Y^*t*^ and pY^*t*^, and results in a reduction of the coefficient of variation of ppY in most of the cases that we considered. Moreover, we also investigated conditions for stochastic dynamic sequestration to have a notable effect, which highly depends on the dynamic range in which the whole system operates. We argued that the time scale of the X protein module must be fast enough such that it can dynamically adapt to changes in the state of the Y protein module, otherwise those changes are averaged out and the correlation in the sample paths disappears. Moreover, the sequestration rate of ppX must have an impact on the X protein module, which is for example not the case if the total number of Y molecules, Y^*T*^, is too small. Depending on operating regimes in the dose response curves of the system, we revealed that dynamic sequestration can also have the opposite effect, namely enhancing output variability. This is the case if the system operates in a regime where ppY is highly sensitive to changes in E_kin_ and at the same time the X protein module is too slow to react instantaneously to state changes in the Y protein module. In this case we observe stochastic oscillations in ppY around its nominal value. The Y protein module reacts sensitively to stochastic changes in ppX, and the response of the X protein module lacks behind. Similar to a negative feedback with a time delay, this leads to oscillating behavior in the state of the Y protein module, and the variation in ppY is increased in this particular case.

So far, retroactive effects have mainly be studied via deterministic approaches. In a recent study (Shah and Vecchio, 2017) it was mathematically shown that retroactivity is attenuated in cascaded phosphorylation and phosphotransfer systems with single and/or double PD cycles with kinase as input, when maintaining a low-high substrate concentration pattern like the MAPK signaling model in Huang and Ferrel (1996). The same architecture with substrate as input is incapable of attenuating retroactivity. Until now, different effects of retroactivity have been described, including the conversion of a graded response into a switch-like response in the context of transcription factor decoy sites (Lee and Maheshri, 2012). Of note, retroactivity via stochastic dynamic sequestration has no direct deterministic counterpart, and it remains a challenging question for the future whether its effect is relevant in real biological systems.

## 4. Materials and Methods

### 4.1. Model representation and assumptions

Models A and B described in the Fig. 1 are expanded using an enzyme-substrate kinetics,

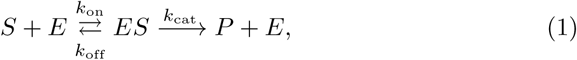

where *S* denoted the substrate. The enzyme *E* is, depending on the particular reaction, either the kinase E_kin_ or the phosphatase E_pho_. Enzyme-substrate complex and product are denoted *ES* and *P*, respectively. Superscripts X and Y refer to the X and the Y protein modules, respectively. In Model B, ppX acts as a kinase for the Y protein. The parameters *k*_on_, *k*_off_ and *k*_cat_ are stochastic rate constants for binding, unbinding and catalytic reactions, respectively. Furthermore, we consider a distributive kinetics for double phosphorylation/dephosphorylation cycles, which requires two separate enzyme-binding events (Salazar and Höfer, 2009).

### 4.2. Parameters and implementation details

**Table 2.** Parameters for Figs 2, 3, 4, 5, 6, 8a, 8c, 8e, and A.9. Stochastic rate constants 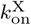 (binding), 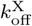 (unbinding) and 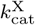 (catalytic) (see Section 4.1 for more details) have units of (molecule^*-*1^time^*-*1^), (time^*-*1^) and (time^*-*1^) respectively.

|  |  |  |  |  |
| --- | --- | --- | --- | --- |
| Initial number of molecules | $X_{\text{init}}$ | $Y_{\text{init}}$ | $E_{\text{pho}}^X$ | $E_{\text{pho}}^Y$ |
|  | 100 | 100 | 20 | 20 |
| Stochastic rate constants | $k_{\text{on}}^X$ | $k_{\text{off}}^X$ | $k_{\text{cat}}^X$ | |
|  | 0.01 | 0.02 | 0.08 |  |

**Table 3.** Parameters for Figs 7, 8b, 8d, and 8f. In Dhananjaneyulu et al. (2012), the values of the Michaelis-Menten (MM) constants of phosphorylation and dephosphorylation reactions for both the X and the Y potein modules are given together with the catalytic rate constants. Here, first a value of *k*_off_ = 0.01 time^*-*1^ is taken, which is within the range of [10^*-*3^ *-* 10^*-*1^] for a typical mammalian cell (Milo, 2013). Subsequently, respective values for *k*on are calculated using the relation 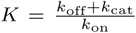, where *K* is the MM constant. For the X system, we denote the MM constants for phosphorylation and dephosphorylation reactions by 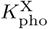 and 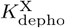, respectively. The same notation applies for the Y system. The values for 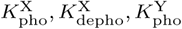, and 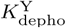 were set to 120, 22, 110, and 22 molecules, respectively, according to Dhananjaneyulu et al. (2012). The corresponding *k*cat values are 0.18 *s*^*-*1^, 0.3 *s*^*-*1^, 0.22*s*^*-*1^ and 0.3 *s*^*-*1^, respectively (Dhananjaneyulu et al., 2012). Values for the range of the number of kinase molecules chosen here, *E*_kin_ = [50, 60, 70, 80, 90] is in the same order of magnitude as the value *E*_kin_ = 94 recorded in Dhananjaneyulu et al. (2012).

|  |  |  |  |  |
| --- | --- | --- | --- | --- |
| Initial number of molecules | $X_{\text{init}}$ | $Y_{\text{init}}$ | $E_{\text{pho}}^X$ | $E_{\text{pho}}^Y$ |
|  | 757 | 567 | 32 | 32 |
| Stochastic rate constants | $k_{\text{on}}^X$ | $k_{\text{off}}^X$ | $k_{\text{on}}^Y$ | $k_{\text{off}}^Y$ |
| **phosphorylation** | 0.0016 | 0.01 | 0.0021 | 0.01 |
| **dephosphorylation** | 0.0141 | 0.01 | 0.0141 | 0.01 |

### 4.3. Numerical simulations

The set of biochemical reactions described by Eq. 1 are simulated using Gillespie’s direct stochastic simulation algorithm implemented in the software *Dizzy* (Ramsey et al., 2005). the expectation values in the steady state and coefficient of variations are obtained from an ensemble of 1000 realizations. To generate individual sample paths, we used our own version of the SSA implemented in MATLAB 2016b (MATLAB, 2016). All simulations were carried out with a laptop equipped with OS X Yosemite Version 10.10.5 having an Intel core i7 processor with a frequency of 2.5 GHz and a memory of 16 GB of type DDR3 with a frequency of 1600 MHz.

## Appendix A Additional figure

**Fig. A.9.**
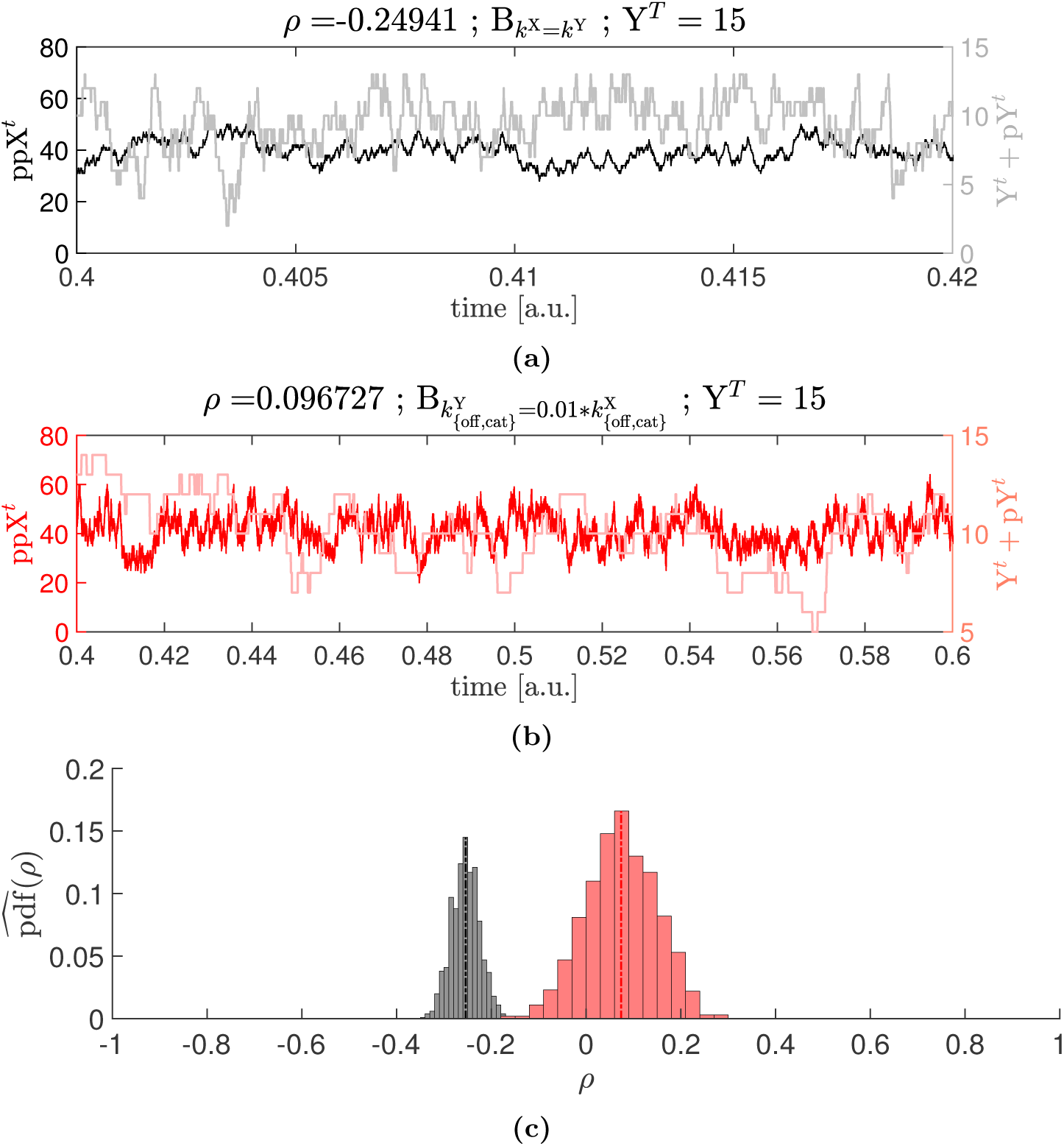
Parameter values: Y^*T*^ = 15, E_kin_ = 21 molecules, *μ*_1_ = *-*0.2544, *σ*_1_ = 0.0297 (black), *μ*_2_ = 0.0739, *σ*_2_ = 0.0745 (red)

## Declarations

### Author’s contributions

DP performed all the simulations. DP and NR designed the study and wrote the main part of the manuscript. All authors discussed the results and implications and commented on the manuscript at all stages of the project. All authors read and approved the final manuscript.

